# SMURF-seq for fast, multiplexed copy number profiling with long-read sequencers

**DOI:** 10.1101/310706

**Authors:** Rishvanth K. Prabakar, Liya Xu, James Hicks, Andrew D. Smith

## Abstract

We present SMURF-seq, a protocol to efficiently sequence short DNA molecules on a long-read sequencer by randomly ligating them to form long molecules. Applying SMURF-seq using Oxford Nanopore Technologies MinION yields up to 30 countable fragments per read at present, which generates multiple copy number profiles in a single run at a reduced time and cost. More broadly, SMURF-seq expands the utility of long-read sequencers for read-counting applications, which do not benefit from increased read length.

---

In the last decade, massively parallel high-throughput short-read sequencing has revolutionized the efficiency and breadth of applications for DNA sequencing [1]. These high-throughput sequencing methods produce millions to billions of short reads in a single run, and have led to the development of many applications that depend on read “counting” to measure the abundance of specific sequences in a sample. These include RNA-seq, ChIP-seq and whole genome copy number profiling. Recently, long-read technologies have been developed that are filling the gap left by short-read sequencers in applications such as genome-assembly [2, 3], which benefit from connecting more distant sequences. Among these the MinION instrument, from Oxford Nanopore Technologies, is highly portable and inexpensive and has shown its unique value for analysis outside of central sequencing facilities [4]. Although long read sequencers such as the MinION typically produce vastly fewer reads from a sequencing run, and are therefore less efficient in applications that use sequenced reads purely as a means to count molecules, they have the enormous advantage of operating in near real-time, with a turnaround time measured in hours rather than days or weeks. We have therefore re-imagined the concatenation sequencing strategy to take advantage of the long-read sequencing format as embodied by the MinION for copy number profiling of cancer samples.

Copy number variation (CNV) has been used successfully to understand a variety of diseases [5] – notably cancers, which exhibit both extreme variation and recurrent trends that can be used for diagnostics and personalized approaches to treatment. For example, the amplification and loss of certain genes, such as RB1 deletion and MYCN amplification in retinoblastoma, can be prognostic or even predictive for treatment [6]. Initial copy number studies used microarray based techniques [7] and were eventually replaced with high-throughput short-read sequencing methods [8]. This led to an increase in the use of CNV profiling for understanding cancer, and also led to profiling single tumor cells [9]. However, the efficiency of high-throughput short-read sequencing for CNV profiles is determined by the availability of instruments and heavy multiplexing of samples. A sequencing core is typically involved and any individual profile must wait for a “full” run before it can be processed. The MinION sequencer has an accessible buy-in, and is easy to use, but has optimal throughput when producing reads that are orders of magnitude longer than needed for CNV profiling.

To make full use of these advantages, we introduce sampling molecules using re-ligated fragments (SMURF)-seq, a protocol to efficiently sequence short DNA molecules on a long-read sequencer for read-counting applications by concatenating short fragments into very long molecules (~8 kb) prior to sequencing. The concept of ligating short molecules before sequencing was introduced in serial analysis of gene expression (SAGE) [10] and subsequently used in short multiply aggregated sequence homologies (SMASH) for CNV profiling using Illumina short read data [11]. SMURF-seq differs from these methods in that (1) the fragmented and re-ligated molecules are longer and the molecule lengths can be highly variable, as suited for long-read sequencing, (2) the protocol involves fewer steps, which in combination with short library preparation time produces sequence data in a few hours, and (3) the sequenced reads are mapped using an approach that tolerates the high error rate of the reads associated with long-read sequencers. Thus SMURF-seq alleviates the issues associated with sequencing short molecules on a long-read sequencer (Supplementary note) and yields sufficient mappable fragments to generate genome-wide CNV profiles on multiple samples in a single run.

For copy number applications described here, the SMURF-seq protocol involves cleaving the genomic DNA into molecules of length that are uniquely mappable to the reference genome. These fragmented molecules are then randomly ligated back together to form artificial, long DNA molecules. The long religated molecules are sequenced following the standard MinION library preparation protocol. SMURF-seq reads are then mapped to the reference genome by splitting them into multiple fragments, each aligning to a distinct region in the genome (Fig. 1). Mapped fragments are grouped into variable length ‘bins’ across the genome and bin counts are used to generate CNV profiles as described in [12].

**Figure 1:**
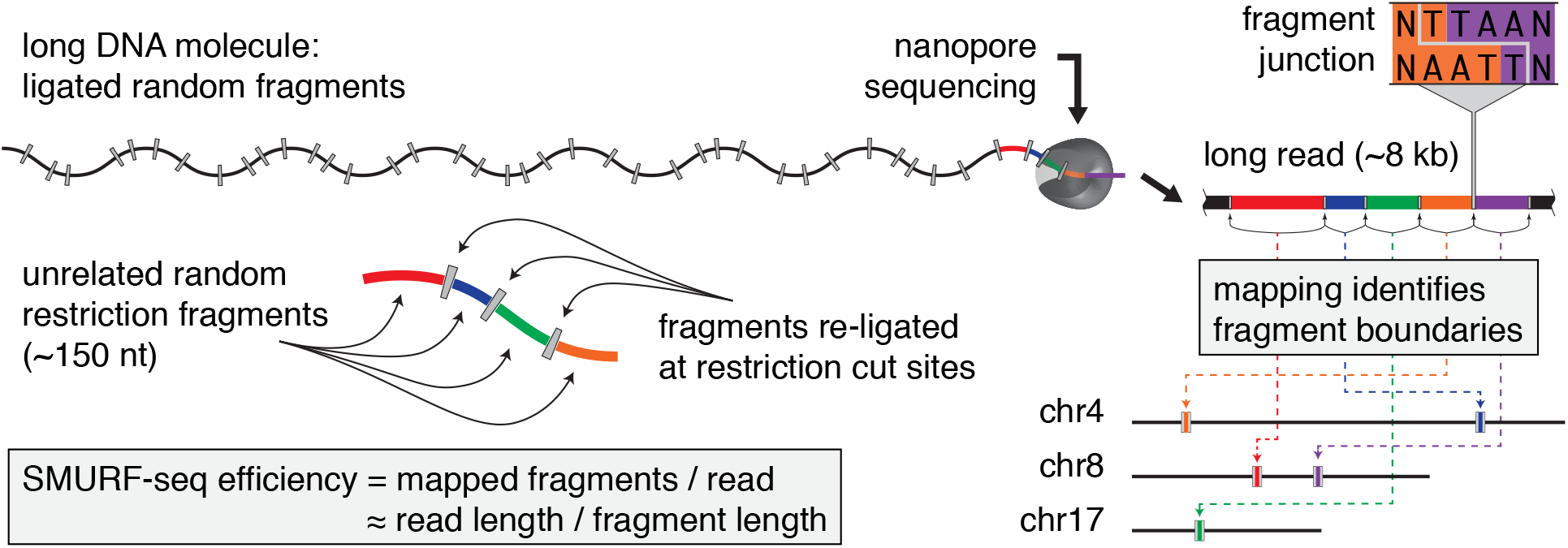
Schematic of SMURF-seq protocol. SMURF-seq efficiently sequences short fragments of DNA for read-counting applications with a reference genome on long-read sequencers, and yields up to 30 countable fragments per sequenced read. SMURF-seq sequences short DNA molecules by generating long concatenated molecules from these. SMURF-seq reads are aligned by splitting them into multiple fragments, each aligning to a distinct region in the genome.

Specifically, the genomic DNA was fragmented using restriction enzymes that have a mean length between restriction sites of approximately 150 bp. We tested SaqAI and Hin1ll restriction enzymes, which produce molecules with mean lengths of 150.2 bp and 208.9 bp respectively (Supplementary Fig. 1). The fragmented DNA molecules were then randomly ligated with T4 DNA ligase enzyme (Supplementary Fig. 1). The resulting long DNA molecules were sequenced following two standard MinION library preparation protocols that are available. SMURF-seq protocol is completely enzymatic and takes less than 90 minutes to complete (Supplementary Fig. 2 and Supplementary Note). We also tested dsDNA Fragmentase enzymes (New England Biolabs) and acoustic shearing (Covaris) to fragment DNA. However, these methods need an additional end-repair step after fragmentation and the ligated molecules were shorter (Supplementary Note).

The reads sequenced using SMURF-seq were mapped to the reference genome by identifying the boundaries of fragments within the reads, done using the seed-and-extend paradigm implemented in BWA-MEM [13]. Briefly, the seed hits from different fragments within a read cluster at different regions of the reference genome. Nearby clusters are joined, and then extended, eventually resulting in one alignment per fragment. BWA-MEM identifies fragment locations with higher than 98.1% precision and 82.6% recall on simulated reads generated by concatenating uniquely mapped fragments (Supplementary Note). These results indicate that (1) existing algorithms, like BWA-MEM, are sufficiently accurate for identifying fragments in SMURF-seq reads, and (2) there is room for improvements in data analysis for SMURF-seq, further improving accuracy of downstream results (e.g. CNV profiles) and total information yield per nucleotide sequenced.

To demonstrate the utility of SMURF-seq, we generated CNV profiles of normal diploid and highly rearranged cancer genomes. Normal diploid female genome was sequenced with SMURF-seq and 267.80k reads (mean read length of 7.1 kb) were generated in one run, which were split into 6.84 million fragments (25.56 mean fragments per read). A flat CNV profile was obtained as expected (Fig. 2a and Supplementary Fig. 3). Then SMURF-seq was applied to a breast cancer line SK-BR-3 and 146.08k reads (mean read length of 8.1 kb) were generated, which were split into 4.24 million fragments (29.03 mean fragments per read). Using a CNV profile obtained by sequencing ~175 million 100 bp paired-end reads from whole genome sequencing (WGS) of SK-BR-3 cells as the gold standard (Supplementary Fig. 9), the profile obtained using SMURF-seq is identical to the gold standard (Pearson r = 0.97; Fig. 2b, c and Supplementary Fig. 4). 5k bins approximately 600 kb in length were used to generate the CNV profiles described above, but higher resolution profiles can be generated using 20k bins as well (Supplementary Fig. 3 and 4). Replicates for normal diploid and SK-BR-3 genomes show a high degree of reproducibility for these profiles (Supplementary Fig. 5 and 6). Normal diploid female genome was also sequenced with SMURF-seq using the Rapid Sequencing Kit (Supplementary Fig. 8), reducing the time to obtain CNV data and demonstrating SMURF-seq with another library preparation protocol.

**Figure 2:**
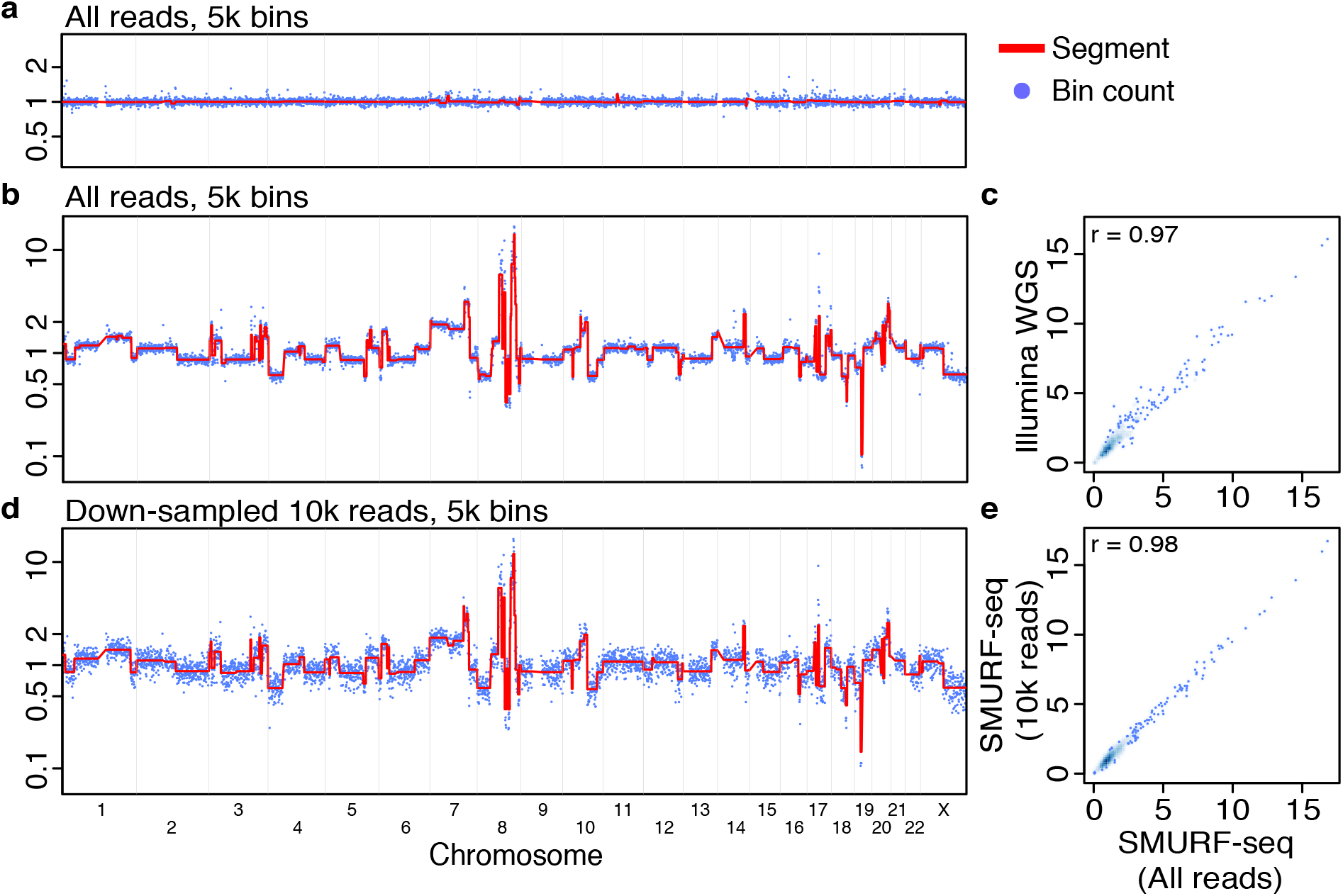
Accurate copy number profiles with SMURF-seq. (a) CNV profile of a normal diploid genome. Each blue point is a normalized bin count and the red line is the segmented bin count. (b) CNV profile of SK-BR-3 cancer genome. (c) Scatter plot of normalized bin counts of SK-BR-3 genome using SMURF-seq and Illumina WGS reads. (d) CNV profile of SK-BR-3 genome with down-sampled 10k SMURF-seq reads. (e) Scatter plot of normalized bin counts of the original SMURF-seq data and data down-sampled to 10k SMURF-seq reads. Pearson correlation of the data is shown.

Using Illumina technology for WGS, 250k reads are sufficient for accurate and quantitative genome-wide CNV profiling [14]. By down-sampling our SMURF-seq data, we verified that 10k reads, approximately 250k fragments, give comparable CNV profiles (Pearson r = 0.98; Fig. 2d and 2e). Thus multiple samples can be barcoded and multiplexed in a single sequencing run. We sequenced two DNA samples (normal diploid female and SK-BR-3) in a single run. After demultiplexing and mapping the reads, the diploid genome had a flat CNV profile as expected and the SK-BR-3 CNV profile is identical to the gold standard (Pearson r = 0.97; Supplementary Fig. 7).

We used SMURF-seq with the low cost MinION sequencer to obtain data similar to that expected from typical short-read sequencing and generated high quality CNV profiles from this output. With a fast and simple preparation method and a turnaround time measured in hours, the SMURF-seq approach could provide a highly efficient methodology for research and clinical laboratories where access to large scale sequencing is limited. CNV profiling in the initial evaluation of cancers has become increasingly important in clinical analysis. At present the MinION instrument has the capacity, using SMURF-seq, to multiplex at least 15 bar-coded samples for CNV profiling in a single run. With continued improvements to the MinION technology this factor will increase, bringing the cost per sample down further.

## Online Methods

### DNA samples

The normal diploid female DNA was purchased from Promega (Cat. no. G1521). Breast cancer cell line SK-BR-3 (American Type of Culture Collection (ATCC), Cat. no. HTB-30) was cultured in RPMI-1640 medium (Thermo Fisher Scientific, Cat. no. 11875093) supplemented with 10% fetal bovine serum (FBS) (Thermo Fisher Scientific, Cat. no. 35011CV) and was maintained at 37° in a humidified chamber supplied with 5% CO2 and was regularly tested for mycoplasma infection.

### Cell lysis and DNA purification

The DNA from SK-BR-3 cells was extracted and purified with the QIAamp DNA Blood Mini Kit (Qiagen, Cat. no. 51104) following the protocol for cultured cells given by the manufacturer. RNA and proteins in the cells were degraded using RNase A stock solution (100 mg/ml) (Qiagen, Cat. no. 19101) and Protease-K (Qiagen, Cat. no. 19133) respectively. Both purchased female diploid DNA and extracted SK-BR-3 DNA were treated with the same downstream processes.

### Fragmenting genomic DNA

2-3 μg of genomic DNA was fragmented with restriction enzyme Anza 64 SaqAI (Thermo Fisher Scientific, Cat. no. IVGN0644) for 30 min at 37°. The fragmented DNA was cleaned with the QIAquick PCR purification kit (Qiagen, Cat. no. 8106) and eluted with 34 μl nuclease-free water. The concentration of DNA was quantified on a Qubit Fluorometer v3 (Thermo Fisher Scientific, cat. no. Q33216) with the Qubit dsDNA HS assay kit (Thermo Fisher Scientific, cat. no. Q32854).

### Ligation of fragmented DNA

500 ng of fragmented DNA in 10 μl nuclease-free water was mixed with 10 μl Anza T4 DNA Ligase Master Mix (Thermo Fisher Scientific, Cat. no. IVGN210-4) and incubated for 30 min at room temperature. The ligated DNA was cleaned with 2 × volume Ampure XP beads (Beckman Coulter, Cat. no. A63881) and eluted in nuclease-free water. This step was done in multiple tubes if more than 500 ng of fragmented DNA was needed to be ligated. The concentration of DNA was quantified on a Qubit Fluorometer v3 with the Qubit dsDNA HS assay kit to ensure ≥1 μg (≥ 400 ng, if the Rapid kit was used for library preparation) remained. The size of the ligated DNA molecules were assessed with 1% agarose gel electrophoresis run at 90 V for 30 min.

### Library preparation (SQK-LSK108 1D DNA by ligation)

1 μg of re-ligated DNA in 45 μl of nuclease-free water was end-repaired and dA-tailed (New England Biolabs (NEB), Cat. no. E7546), followed by elution in nuclease-free water after 1.5 × volume Ampure XP beads clean-up. Sequencing adapters (AMX1D) were ligated with Blunt/TA Ligase Master Mix (NEB, Cat.no. M0367) and cleaned with 0.4× volume Ampure XP beads and eluted using 15 μl Elution Buffer (ELB) following the manufacture’s protocol (Oxford Nanopore Technologies (ONT), 1D genomic DNA by ligation protocol).

### Multiplexed library preparation (EXP-NBD103 and SQK-LSK108)

700 ng of each re-ligated sample in 45 μl of nuclease-free water was end-repaired and dA-tailed (NEB, Cat. no. E7546) and cleaned with 1.5 × volume Ampure XP beads and eluted in nuclease-free water. Different Native Barcodes (NB-x) for each sample was ligated with Blunt/TA Ligase Master Mix (NEB, Cat.no. M0367) and cleaned with 2× volume Ampure XP beads and eluted in nuclease-free water. Equimolar amounts of each sample was pooled to have 700 ng of DNA in 50 μl water. Barcode adapters (BAM) was ligated with Quick T4 DNA Ligase (NEB, Cat. no. E6056) and cleaned with 0.4 × volume Ampure XP beads and eluted using 15 μl Elution Buffer (ELB) following the manufacture’s protocol (ONT, 1D native barcoding genomic DNA).

### Library preparation (SQK-RAD003 Rapid sequencing)

400 ng of re-ligated DNA was concentrated with 2 × volume Ampure XP beads to 7.5 μl nuclease-free water. DNA was tagmented with Fragmentation Mix (FRA), and Rapid 1D Adapter (RPD) was attached following the manufacture’s protocol (ONT, rapid sequencing).

### MinION sequencing and base-calling

All the prepared libraries were loaded on R9.5 Flowcells following the manufacture’s protocol (ONT) and sequenced for up to 48 hours using the script specific to library preparation protocol. Base-calling and de-multiplexing barcoded reads were performed using ONT Albacore Sequencing Pipeline Software (1.2.6) with the appropriate parameters based on the library preparation kit.

### Read alignment

The sequenced reads were mapped to the human reference genome (hg19) with BWA-MEM (0.7.17) with the “-x ont2d -k 1 -W 5” options (Supplementary Note).

### Estimation of copy number variations

CNV profiles were generated using the pipeline provided in the supplementary of [12]. The human reference genome (hg19) was split into 5k (or 20k) bins containing the same number of uniquely mappable locations. The counts for each bin was determined from the uniquely mapped fragments, and normalized for GC content using LOWESS smoothing. Bins with spuriously high bin counts (“bad bins”) were removed. Circular binary segmentation (CBS) [15] was used to identify breakpoints in the normalized bin counts.

### Comparison with published SK-BR-3 data

SK-BR-3 genome sequenced from a population of cells using 100 bp paired-end reads was downloaded from SRA (SRR504589) [16]. The reads were mapped with BWA-MEM using the default parameters, and CNV profiles were generated using the uniquely mapped and properly paired reads. The scatter plots and the Pearson correlations comparing the CNV profiles were produced using R.

## References

[1] Martin Kircher and Janet Kelso. High-throughput dna sequencing–concepts and limitations. Bioessays, 32(6):524–536, 2010.

[2] Miten Jain, Sergey Koren, Karen H Miga, Josh Quick, Arthur C Rand, Thomas A Sasani, John R Tyson, Andrew D Beggs, Alexander T Dilthey, Ian T Fiddes, et al. Nanopore sequencing and assembly of a human genome with ultra-long reads. Nature biotechnology, 2018.

[3] Nicholas J Loman, Joshua Quick, and Jared T Simpson. A complete bacterial genome assembled de novo using only nanopore sequencing data. Nature methods, 12(8):733, 2015.

[4] Joshua Quick, Nicholas J Loman, Sophie Duraffour, Jared T Simpson, Ettore Severi, Lauren Cowley, Joseph Akoi Bore, Raymond Koundouno, Gytis Dudas, Amy Mikhail, et al. Real-time, portable genome sequencing for ebola surveillance. Nature, 530(7589):228, 2016.

[5] Jonathan Sebat, B Lakshmi, Dheeraj Malhotra, Jennifer Troge, Christa Lese-Martin, Tom Walsh, Boris Yamrom, Seungtai Yoon, Alex Krasnitz, Jude Kendall, et al. Strong association of de novo copy number mutations with autism. Science, 316(5823):445–449, 2007.

[6] Jesse L Berry, Liya Xu, A Linn Murphree, Subramanian Krishnan, Kevin Stachelek, Emily Zolfaghari, Kathleen McGovern, Thomas C Lee, Anders Carlsson, Peter Kuhn, et al. Potential of aqueous humor as a surrogate tumor biopsy for retinoblastoma. JAMA ophthalmology, 135(11):1221–1230, 2017.

[7] Robert Lucito, John Healy, Joan Alexander, Andrew Reiner, Diane Esposito, Maoyen Chi, Linda Rodgers, Amy Brady, Jonathan Sebat, Jennifer Troge, et al. Representational oligonucleotide microarray analysis: a high-resolution method to detect genome copy number variation. Genome research, 13(10):2291–2305, 2003.

[8] Derek Y Chiang, Gad Getz, David B Jaffe, Michael JT O’kelly, Xiaojun Zhao, Scott L Carter, Carsten Russ, Chad Nusbaum, Matthew Meyerson, and Eric S Lander. High-resolution mapping of copy-number alterations with massively parallel sequencing. Nature methods, 6(1):99, 2009.

[9] Nicholas Navin, Jude Kendall, Jennifer Troge, Peter Andrews, Linda Rodgers, Jeanne McIndoo, Kerry Cook, Asya Stepansky, Dan Levy, Diane Esposito, et al. Tumour evolution inferred by single-cell sequencing. Nature, 472(7341):90, 2011.

[10] Victor E Velculescu, Lin Zhang, Bert Vogelstein, and Kenneth W Kinzler. Serial analysis of gene expression. Science, 270(5235):484–487, 1995.

[11] Zihua Wang, Peter Andrews, Jude Kendall, Beicong Ma, Inessa Hakker, Linda Rodgers, Michael Ronemus, Michael Wigler, and Dan Levy. Smash, a fragmentation and sequencing method for genomic copy number analysis. Genome research, 26(6):844–851, 2016.

[12] Timour Baslan, Jude Kendall, Linda Rodgers, Hilary Cox, Mike Riggs, Asya Stepansky, Jennifer Troge, Kandasamy Ravi, Diane Esposito, B Lakshmi, et al. Genome-wide copy number analysis of single cells. Nature protocols, 7(6):1024, 2012.

[13] Heng Li. Aligning sequence reads, clone sequences and assembly contigs with bwa-mem. arXiv preprint arXiv:1303.3997, 2013.

[14] Timour Baslan, Jude Kendall, Brian Ward, Hilary Cox, Anthony Leotta, Linda Rodgers, Michael Riggs, Sean D’Italia, Guoli Sun, Mao Yong, et al. Optimizing sparse sequencing of single cells for highly multiplex copy number profiling. Genome research, 25(5):714–724, 2015.

[15] Adam B Olshen, ES Venkatraman, Robert Lucito, and Michael Wigler. Circular binary segmentation for the analysis of array-based dna copy number data. Biostatistics, 5(4):557–572, 2004.

[16] Yong Wang, Jill Waters, Marco L Leung, Anna Unruh, Whijae Roh, Xiuqing Shi, Ken Chen, Paul Scheet, Selina Vattathil, Han Liang, et al. Clonal evolution in breast cancer revealed by single nucleus genome sequencing. Nature, 512(7513):155, 2014.

